# OpenLabCluster: Active Learning Based Clustering and Classification of Animal Behaviors in Videos Based on Automatically Extracted Kinematic Body Keypoints

**DOI:** 10.1101/2022.10.10.511660

**Authors:** Jingyuan Li, Moishe Keselman, Eli Shlizerman

**Affiliations:** Department of Electrical and Computer Engineering, University of Washington, Seattle, WA; Department of Applied Mathematics, University of Washington, Seattle, WA

## Abstract

Quantifying natural behavior from video recordings is a key component in ethological studies. Markerless pose estimation methods have provided an important step toward that goal by automatically inferring kinematic body keypoints. Such methodologies warrant efficient organization and interpretation of keypoints sequences into behavioral categories. Existing approaches for behavioral interpretation often overlook the importance of representative samples in learning behavioral classifiers. Consequently, they either require extensive human annotations to train a classifier or rely on a limited set of annotations, resulting in suboptimal performance. In this work, we introduce a general toolset which reduces the required human annotations and is applicable to various animal species. In particular, we introduce OpenLabCluster, which clusters temporal keypoint segments into clusters in the latent space, and then employ an Active Learning (AL) approach that refines the clusters and classifies them into behavioral states. The AL approach selects representative examples of segments to be annotated such that the annotation informs clustering and classification of all temporal segments. With these methodologies, OpenLabCluster contributes to faster and more accurate organization of behavioral segments with only a sparse number of them being annotated. We demonstrate OpenLabCluster performance on four different datasets, which include different animal species exhibiting natural behaviors, and show that it boosts clustering and classification compared to existing methods, even when all segments have been annotated. OpenLabCluster has been developed as an open-source interactive graphic interface which includes all necessary functions to perform clustering and classification, informs the scientist of the outcomes in each step, and incorporates the choices made by the scientist in further steps.

## 1 Introduction

Analysis and interpretation of animal behavior are essential for a multitude of biological investigations. Behavioral studies extend from ethological studies to behavioral essays as a means to investigate biological mechanisms [1, 3, 4, 5, 6, 7, 8, 9, 10, 2]. In these studies, methodologies facilitating robust, uninterrupted, and high-resolution observations are key. Indeed, researchers have been recording animal behaviors for decades with various modalities, such as video, sound, placement of physical markers, and more [11, 12, 13, 14, 15, 16, 17]. Recent enhancements in recording technologies have extended the ability for the deployment of recording devices in various environments and for extended periods of time. The enhancement in the ability to perform longer observations and in the number of modalities brings forward the need to organize, interpret, and associate recordings with identified repertoires of behaviors, i.e., perform classification of the recordings into behavioral states. Performing these operations manually would typically consume a significant amount of time and would require expertise. For many recordings, manual behavior classification becomes an unattainable task. Therefore, it is critical to develop methodologies to accelerate the classification of behavioral states and require as little involvement from the empiricist as possible [18, 19, 20].

Early efforts in automatic behavior classification focused on raw video analysis using machine learning techniques such as Convolutional Neural Networks (CNNs) [22, 23, 24, 25], Recurrent Neural Networks (RNNs) [26, 27], Temporal Gaussian Mixture models [28], and temporal CNNs [29]. While effective in specific scenarios, video-based methods often incorporate extraneous background information and noise (e.g., camera artifacts), which can undermine reliability and require considerable computational resources due to the high-dimensional nature of video data [29]. In contrast, approaches that concentrate on movement by utilizing body keypoints or kinematics—extracted from video frames—can circumvent these limitations [30, 31, 32, 33, 34].

Markerless pose estimation techniques, such as OpenPose, DeepLabCut, Anipose, and others [35, 36, 37, 38, 39, 40], enable accurate keypoint detection without the need for physical markers. Furthermore, numerous related tools and approaches have also been further introduced to advance animal pose estimation [2, 41, 42, 43, 44, 45, 46]. Once body keypoints are estimated, behavioral segmentation can be achieved using unsupervised clustering methods—such as HBDSCAN [47, 48], hierarchical clustering [49], and the Watershed algorithm [50, 51, 52]—which group similar postural states and differentiate distinct behaviors [4, 48]. Dimensionality reduction techniques, including Principal Component Analysis (PCA) and Uniform Manifold Approximation and Projection (UMAP), further enhance the representation of body keypoints for effective clustering [53, 49, 54, 48, 55].

Recent deep learning methods have advanced latent keypoint representation learning through task-specific optimization, as demonstrated by TREBA [56] and its automated extension, AutoSWAP [57]. Contrastive learning approaches have also been proposed to refine the latent space by drawing together similar behavioral samples and separating dissimilar ones [58, 59]. One of the challenges in such approaches is the selection of appropriate positive and negative samples remains challenging without human guidance. General methods such as Predict&Cluster [60] and VAME [61] address these challenges by focusing on sequence reconstruction and future prediction, enabling the unsupervised clustering of behavioral patterns [60, 61, 62].

While unsupervised clustering can identify similar behavioral patterns [60, 61, 62], it may not effectively identify behaviors of specific interest. Supervised classification approaches address this limitation by mapping behavioral segments to behavioral categories of interest, under the guidance of annotated training data [30, 33, 63]. The classification accuracy critically depends on both the choice of classifier and the quality and quantity of annotations. Early success in behavioral classification was achieved using classical machine learning approaches [64, 21, 63, 65, 66, 67, 68, 48]. These have recently been supplemented by deep learning approaches[69, 70, 58, 46, 33]. Nevertheless, manual annotation remains labor-intensive and subject to inter-annotator variability.

To address the need for manual annotation, methods such as SaLSa—which assigns uniform labels to pre-computed unsupervised clusters—and JAABA—which provides an interactive framework for correcting misclassifications—have been developed [70, 71]. Active learning (AL) techniques further streamline the process by automatically selecting samples for annotation, balancing annotation effort with classification accuracy [72, 73, 74, 75, 76]. In particular, for behavior recognition, A-SOiD employs AL to prioritize samples with high prediction uncertainty for animal behavior recognition; however, uncertainty-based selection may inadvertently target redundant samples. Integrating clustering information with classifier uncertainty could improve the efficiency of sample selection.

In this work, we extend previous methods by jointly learning representations for AL and classifier training using pose estimated from video recordings, e.g., keypoints estimated using DeepLabCut [35, 38, 36]. In particular, we introduce the OpenLabCluster toolset, AL based semi-supervised behavior classification platform embedded in a graphic interface for animal behavior classification from body keypoints. The system implements and allows the use of multiple semi-supervised AL methods. AL is performed in an iterative way, where, in each iteration, an automatic selection of a subset of candidate segments is chosen for annotation, which in turn enhances the accuracy of clustering and classification. OpenLabCluster is composed of two components illustrated in Fig. 1A: (1) Unsupervised deep encoder-decoder clustering of behavior representation, *Cluster Maps*, which depict the representations as points and show their groupings, followed by (2) Iterative automatic selection of representations for annotation and subsequent generation of *Behavior Classification Maps*. In each iteration, each point in the Cluster Map is re-positioned and associated with a behavioral class (colored with a color that corresponds to a particular class). These operations are performed through the training of a clustering encoder-decoder (component (1)) along with a deep classifier (component (2)). OpenLabCluster implements these methodologies as an open-source graphical user interface (GUI) to empower scientists with little or no deep-learning expertise to perform animal behavior classification. In addition, OpenLabCluster includes advanced options for experts.

**Figure 1.**
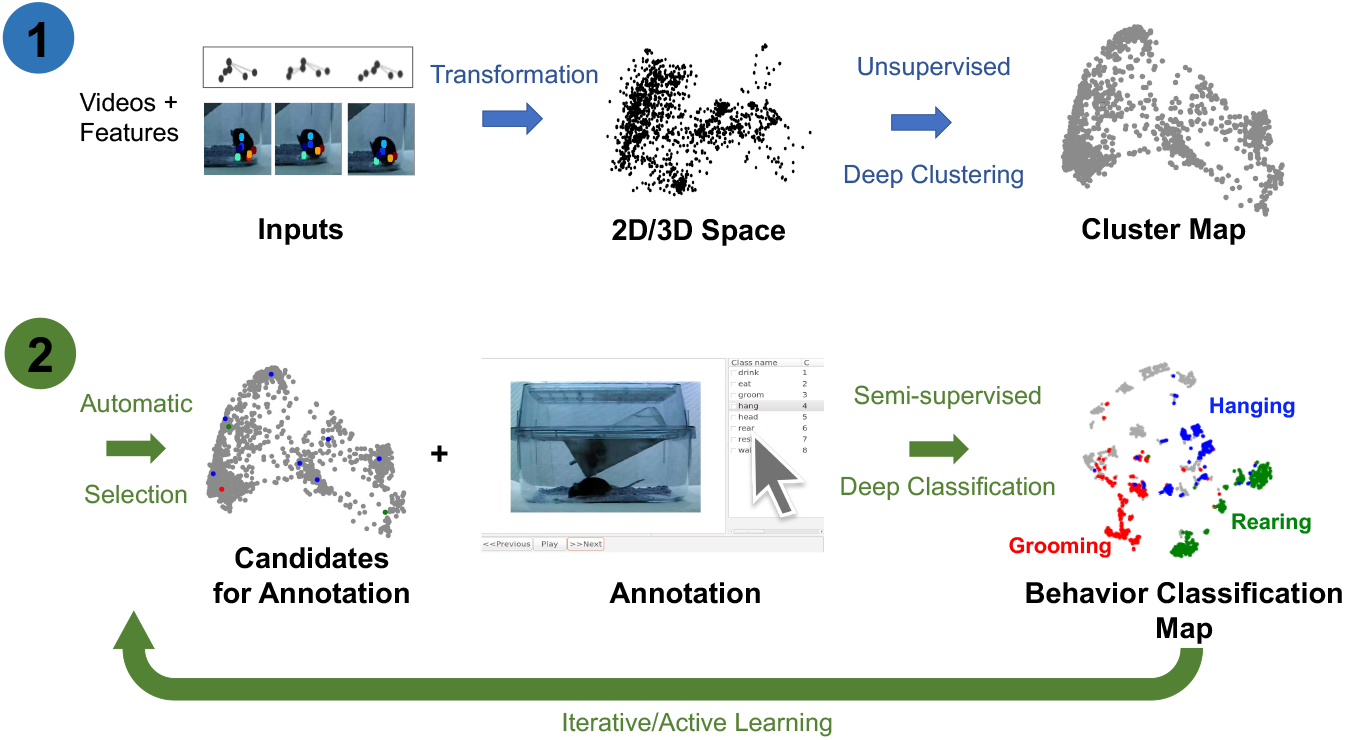
OpenLabCluster overview. (1) Clustering: Input of body keypoints segments is mapped to low dimensional space. Unsupervised encoder-decoder maps them to a Cluster Map. (2) Classification: AL algorithms automatically select candidates for annotation after which Cluster Map is reshaped into the Behavior Classification Map where each point is associated with a behavioral state (Grooming (red), Hanging (blue), Rearing (blue)). Mouse images are reproduced from frames of videos provided in the dataset of [21].

## 2 Results

### Datasets

Behavioral states and their dynamics vary from species to species and from recordings to recordings. We use four different datasets to demonstrate OpenLabCluster applicability to various settings. The datasets include videos of behaviors of four different animal species (Mouse [21], Zebrafish [4], *C. elegans* [77], Monkey [42]) with three types of motion features (body keypoints, kinematics, segments), as depicted in Fig. 2. Two of the datasets include apriori annotated behavioral states (ground truth) (Mouse, *C. elegans*), while the Zebrafish dataset includes ground truth a priori predicted by another method, and the Monkey dataset does not include ground truth annotations. Three of the datasets have been temporally segmented into *single-action clips* (Mouse, Zebrafish, *C. elegans*), i.e., temporal segments while the Monkey dataset is a continuous recording that requires segmentation into clips. We describe further details about each dataset below.

**Figure 2.**
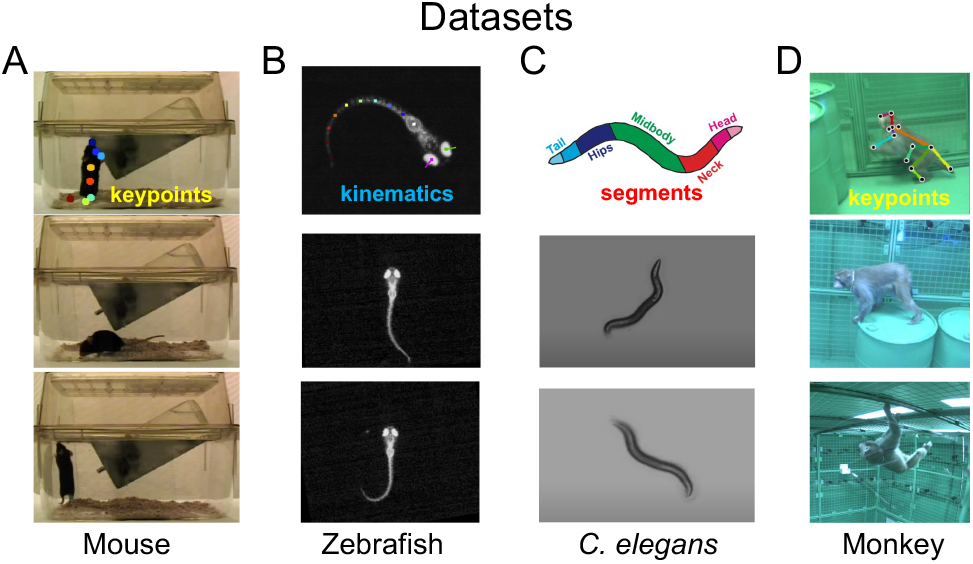
Visualization of four animal behavior datasets. (A) Home-Cage mouse dataset (Mouse) (B) *C. elegans* Movement Dataset (*C. elegans*) (C) Zebrafish free swimming dataset (Zebrafish) (D) OpenMonkeyStudio Macaque behaviors dataset (Monkey). The top row shows positions of extracted keypoints for each dataset. Images sources: A. Images are reproduced from frames of videos in the dataset of [21]. B. Images are reproduced from Figure 1 and frames of videos of [4]. C. Images are reproduced from Supplementary Figure 1 and images of [77]. D. Images are reproduced from Figure 8 and images of [42].

#### Home-Cage Mouse

The dataset includes video segments of 8 identified behavioral states [21]. In particular, it contains videos recorded by front cage cameras when the mouse is moving freely and exhibits natural behaviors, such as drinking, eating, grooming, hanging, micromovement, rearing, walking, and resting. Since keypoints have not been provided in this dataset, we use DeepLabcut [35, 38, 36] to automatically mark and track eight body joint keypoints (snout, left-forelimb, right-forelimb, left-hindlimb, right-hindlimb, fore-body, hind-body, and tail) in all recorded videos frames. An example of estimated keypoints overlaid on top of the corresponding video frame is shown in Fig. 2A (top). To reduce the noise that could be induced by the pose estimation procedure, we only use the segments for which DeepLabCut estimation confidence is high enough. We use 8 sessions for training the models of clustering and classification (2856 segments) and test classification accuracy on 4 other sessions (393 segments).

#### Zebrafish

The dataset includes video footage of zebrafish movements and was utilized in [4] for unsupervised behavior clustering using 101 precomputed kinematic features, a procedure that identified 13 clusters which were manually related to 13 behavior prototypes (see Appendix 5.2). In the application of OpenLabCluster to this dataset, we utilize only a small subset of these features (16 features) and examine whether OpenLabCluster is able to generate classes aligned with the unsupervised clustering results obtained on full 101 features (as the ground truth). We use 5294 segments for training and 2781 segments for testing.

#### C. elegans

The dataset is recorded with Worm Tracker 2.0 when the worm is freely moving. The body contour is identified automatically using contrast to background from which kinematic features are calculated and constitute 98 features that correspond to body segments from head to tail in 2D coordinates, see [77] and Fig. 2C. Behavioral states are divided into three classes: moving forward, moving backward, and staying stationary. We use ten sessions (a subset) to investigate the application of OpenLabCluster to this dataset, where the first 7 sessions (543 segments) are used for training and the remaining 3 sessions (196 segments) are used for testing.

#### Monkey

This dataset is from OpenMonkeyStudio repository [42] and captures freely moving macaques in a large unconstrained environment using 64 cameras encircling an open enclosure. 3D keypoints positions are reconstructed from 2D images by applying deep neural network reconstruction algorithms on the multi-view images. Among the movements, 6 behavioral classes have been identified. In contrast to other datasets, this dataset consists of continuous recordings without segmentation into action clips. We thereby segment the videos by clipping them into fixed duration clips (10 frames with 30 fps rate) which results in 919 segments, where each segment is ≈ 0.33 seconds long. OpenLabCluster receives the 3D body key points of each segment as inputs. Notably, a more advanced technology could be implemented to segment the videos as described in [78]. Here, we focused on examining the ability of OpenLabCluster to work with segments that have not been pre-analyzed and thus used the simplest and most direct segmentation method.

### Evaluation Metrics

We evaluate the accuracy of OpenLabCluster by computing the percentage of temporal segments in the test set that OpenLabCluster correctly associated with the states given as ground truth, such that 100% accuracy will indicate that OpenLabCluster correctly classified all temporal segments in the test set. Since OpenLabCluster implements the semi-supervised approach to minimize the number of annotations for segments, we compute the accuracy given annotation budgets of overall 5%, 10%, 20% labels to be used over the possible iterations in conjunction with AL. In particular, we test the accuracy when the Top, CS, and MI AL methods implemented in OpenLabCluster are used for the selection of temporal segments to annotate.

### Benchmark Comparison

We evaluated OpenLabCluster against established animal behavior classification approaches either with or without AL methods. For non-AL approaches, we compared against K-Nearest Neighbour (KNN) [79], Support Vector Machine (SVM) [21], SimBA’s Random Forest Classifier (RFC) [68], and VAME with an additional classifier (VAME+C) [61]. With respect to AL, we compared our method to A-SOiD [76], which employs RFC with selective sampling. Furthermore, we conducted ablation studies by evaluating OpenLabCluster with decoder removed and explored alternative architectures by integrating VAME’s encoder-decoder (OpenLabCluster-V) with various AL strategies (CS, TOP, and MI). Detailed experimental settings and additional results are provided in the Benchmark Details Section 4.

### Outcomes

The results of evaluation in 5 runs are shown in Tables 1 and 2 and further analysis in Fig. 3 and Fig. 4. We summarize the main outcomes of the evaluation and their interpretation below.

**Table 1:** Classification accuracy of Home-Cage Mouse behaviors for increasing number of annotated segments. Top: Classification accuracy of existing baselines: KNN, SVM, C, SimBA, A-SOiD, and VAME+C. Middle: Accuracy of OpenLabCluster using three AL strategies (CS, Top, and MI). Bottom: Accuracy of OpenLabCluster with the VAME encoder-decoder (OpenLabCluster-V). Boldface indicates the best accuracy.

| Mouse (8 classes; Keypoints) |  |  |  |  |
| --- | --- | --- | --- | --- |
| Labels (%) | 5 | 10 | 20 | 100 |
| Labels (#) | 143 | 286 | 571 | 2856 |
| KNN [79] | 43.5±3.9 | 53.1±1.7 | 51.5±2.9 | 60.8±0.0 |
| SVM[21] | 50.6±6.0 | 60.3±2.6 | 64.6±1.6 | 72.3±0.0 |
| C | 55.2±3.6 | 60.7±1.7 | 64.5±2.1 | 71.5±1.0 |
| SimBA[68] | 62.2±3.8 | 66.5±2.2 | 69.8±1.2 | 79.8±0.9 |
| A-SOiD[76] | 63.7±3.9 | 60.1±3.4 | 65.5±1.7 | 70.9±0.9 |
| VAME+C[61] | 67.4±4.6 | 75.1±1.8 | 77.8±1.3 | 85.2±0.7 |
| OpenLabCluster Top | 58.6±2.6 | 69.2±1.8 | 76.7±1.2 | 83.8±0.4 |
| OpenLabCluster MI | 65.8±2.8 | 76.6±0.9 | 79.1±0.7 | 82.0±0.7 |
| OpenLabCluster CS | 66.2±3.1 | 74.5±1.8 | **81.5±1.0** | 83.9±0.3 |
| OpenLabCluster-V TOP | 68.3±1.8 | 75.3±1.8 | 77.5±1.2 | 85.6±0.8 |
| OpenLabCluster-V CS | 74.4±1.4 | **77.7±1.8** | 81.4±0.9 | **85.8±0.7** |
| OpenLabCluster-V MI | **74.7±1.2** | 76.1±1.5 | 80.1±0.6 | 85.3±0.3 |

**Table 2:** Classification accuracy of Zebrafish and *C. elegans* behaviors for increasing the number of annotated segments (reported as percentage (%)). Top: benchmark methods KNN, SVM, and C, SimBA, A-SOiD, VAME+C, C. Middle: Accuracy of OpenLabCluster using various AL approaches: CS, Top, and MI. Bottom: OpenLabCluster with VAME encoder-decoder (OpenLabCluster-V). Best accuracy is highlighted in boldface.

|  | Zebrafish (13 classes; Kinematics) |  |  |  | *C. elegans* (3 classes; Segments) |  |  |  |
| --- | --- | --- | --- | --- | --- | --- | --- | --- |
| Labels (%) | 5 | 10 | 20 | 100 | 5 | 10 | 20 | 100 |
| Labels (#) | 265 | 530 | 1059 | 5294 | 27 | 55 | 109 | 543 |
| SVM[21] | 55.8±1.5 | 63.9±1.2 | 68.6±0.5 | 74.0±0.0 | 76.3±0.8 | 76.4±0.3 | 74.5±0.6 | 76.0±0.0 |
| KNN [79] | 57.2±1.0 | 60.3±1.1 | 63.0±0.3 | 69.2±0.0 | 61.0±11.3 | 71.5±3.4 | 73.3±2.6 | 77.0±0.0 |
| VAME+C[61] | 62.1±1.1 | 67.7±0.4 | 70.9±0.5 | 75.2±0.3 | 73.8±3.8 | 75.7±1.5 | 76.6±0.2 | 76.5±0.0 |
| C | 65.7±1.6 | 70.0±0.7 | 75.8±0.9 | 82.8±0.1 | 71.3±4.7 | 75.2±3.4 | 77.5±0.9 | **94.8±0.7** |
| A-SOiD[76] | 68.7±0.7 | 72.3±0.5 | 74.5±0.4 | 77.8±0.1 | 73.4±5.6 | 74.4±2.5 | 76.6±1.0 | 80.1±1.0 |
| SimBA[68] | 69.7±1.0 | 73.0±0.8 | 75.9±0.7 | 80.3±0.3 | 68.9±12.7 | 74.1±4.1 | 77.2±2.1 | 83.5±0.6 |
| OpenLabCluster CS | 71.9±1.0 | 76.6±1.0 | 79.8±0.6 | 83.2±0.1 | 73.5±1.0 | 64.3±5.1 | 75.1±3.1 | 92.8±0.4 |
| OpenLabCluster Top | 72.0±1.2 | 77.1±0.7 | 80.1±0.7 | **83.5±0.4** | 76.5±0.0 | 76.5±3.1 | **77.9±1.1** | 93.0±0.5 |
| OpenLabCluster MI | **74.7±0.7** | **79.2±0.3** | **81.1±0.4** | 83.4±0.2 | 76.6±0.2 | 74.7±3.0 | 76.9±0.2 | 93.6±0.8 |
| OpenLabCluster-V CS | 44.4±1.6 | 53.5±2.6 | 64.3±1.3 | 75.1±0.2 | 65.6±12.6 | 76.5±0.3 | 75.4±2.4 | 76.6±0.2 |
| OpenLabCluster-V TOP | 63.0±1.0 | 67.8±0.8 | 71.6±0.9 | 75.1±0.2 | **75.8±2.2** | **76.8±0.2** | 76.7±0.2 | 76.5±0.0 |
| OpenLabCluster-V MI | 65.9±0.7 | 69.4±0.7 | 71.9±0.4 | 75.1±0.3 | 75.7±1.2 | 76.8±0.4 | 76.8±0.4 | 76.5±0.0 |

**Figure 3.**
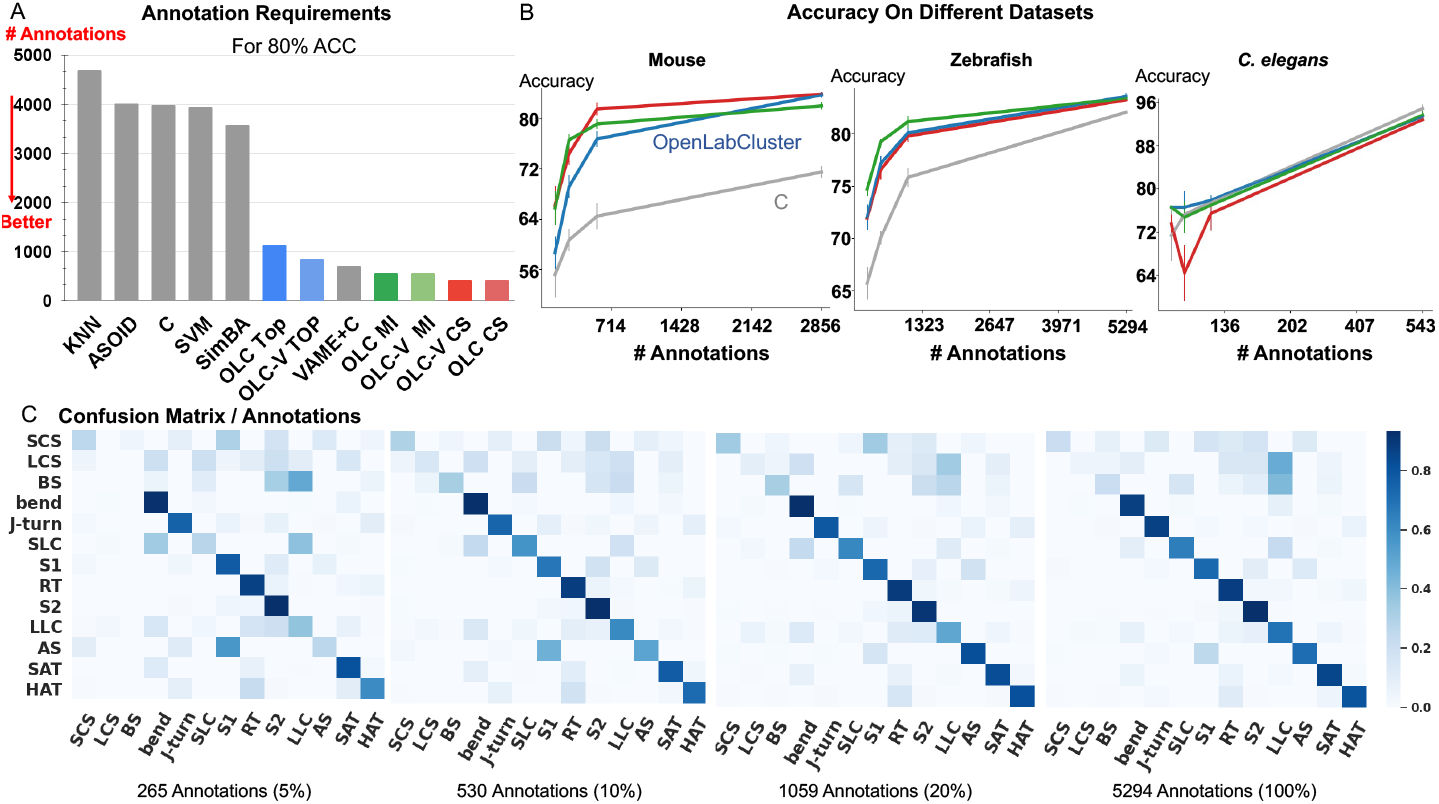
Relation Between Accuracy and Annotation. A: The amount of annotations required to achieve 80% accuracy for classification of Home-Cage Mouse behaviors. Computed for benchmark methods (KNN, SVM, and C, SimBA, A-SOiD, VAME+C), and variants of OpenLabCluster with three AL methods (Top, MI, CS). B: Prediction accuracy with increasing annotation budget on three datasets of Mouse, *C. elegans* and Zebrafish. C: Confusion matrix for zebrafish dataset for increasing annotation budget (5%, 10%, 20%, 100%).

**Figure 4.**
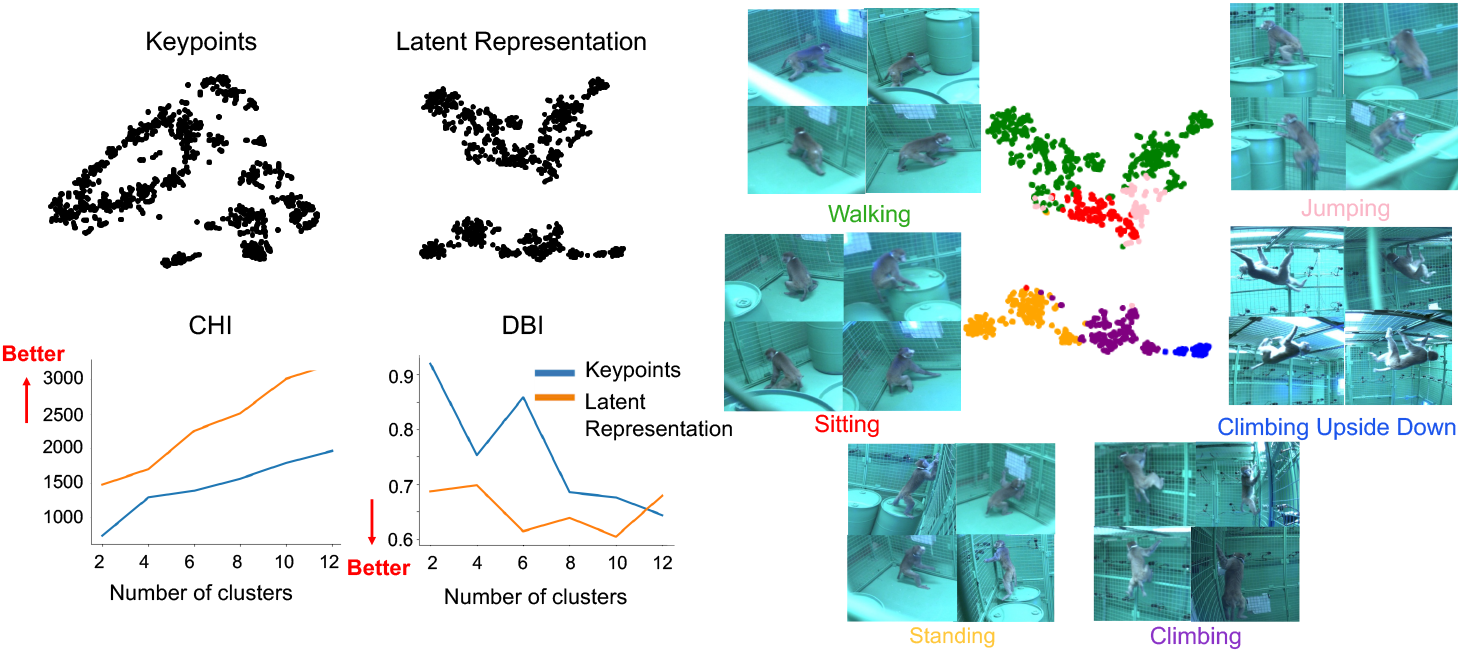
2D tSNE projection of behavioral segments. Left, Top: 2D tSNE projection of keypoints compared with 2D tSNE projection of Latent Representation (Cluster Map). Left, Bottom: CHI and DBI metrics computed for each projection for increasing number of clusters. Right: Behavior Classification Map (2D tSNE projection of Latent Representation with associated behavioral states colors) along with example video frames from segments associated with each class.Images are reproduced from images in the datset of [42].

#### Accuracy of Classification

We observe that *the accuracy of classification of OpenLabCluster across datasets is consistently higher than standard supervised classification methods (e*.*g*., *C, SVM, SimBA) for almost any budget of annotation*. Specifically, for the Home-Cage Mouse Behavior dataset Table 1, OpenLabCluster achieves the accuracy of 66.2% when just 143 (5% of 2856) segments have been annotated. Accuracy improves along with the increase in the number of annotated segments, i.e., accuracy is 76.6% when 10% of segments are annotated, and 81.5% when 20% of segments are annotated. Compared to C—the encoder-only version of OpenLabCluster—OpenLabCluster achieves an average accuracy increase of approximately 12%. This improvement underscores the importance of both the encoder-decoder structure and active sample selection. Meanwhile, VAME+C, which incorporates an encoder-decoder and a classifier, shows promise by reaching an accuracy of 67.4% with only 5% annotated samples. However, it remains less optimal than OpenLabCluster-V, which attains 74.7% under the same annotation budget. These results further illustrate the effectiveness of AL for accurate behavior classification with limited labels. It should be noted that although A-SOiD also integrates active learning for sample selection, its classifier design and exclusive reliance on an uncertainty-based selection method appears to limit its performance relative to OpenLabCluster variants. Among the AL strategies TOP, CS, and MI, both CS and MI outperform TOP on the Home-Cage Mouse dataset. Notably, while AL is expected to be especially effective in sparse annotation scenarios when all segments are annotated (fully supervised scenario) the accuracy of the OpenLabCluster MI approach exceeds supervised classification approaches (C) by 12.4% (rightmost column in Table 1). This reflects the effectiveness of the targeted selection of candidates for annotation and the use of clustering latent representation to enhance the overall organization of the segments.

For Zebrafish and *C. elegans* datasets, OpenLabCluster consistently achieves higher accuracy, except when 100% of the *C. elegans* dataset is annotated, demonstrating its generalizability across various animal behavior datasets. When compared to its encoder-only variant (C), OpenLabCluster exhibits an accuracy improvement of approximately 7.8% on the zebrafish dataset and around 2.2% on the *C. elegans* dataset. These gains are less pronounced than those observed on the Home-Cage Mouse dataset. These could be associated with not having manually identified ground truth behavior states for Zebrafish and having only 3 classes for the *C. elegans* dataset which is a simpler semantic task that does not challenge classifiers. We can indeed observe that when all annotations are considered in *C. elegans* dataset, all approaches perform well (above 92%) and a standard classifier achieves the best accuracy. In contrast to the results observed with the Home-Cage Mouse dataset, we find that on the Zebrafish dataset, OpenLabCluster-V underperforms OpenLabCluster and exhibits comparable performance on *C. elegans*. Moreover, the CS strategy appears unsuitable for OpenLabCluster-V in the Zebrafish setting, resulting in diminished outcomes.

#### The Amount of Required Annotations

Since accuracy varies across datasets and depends on the number of classes and other aspects, we examine the relationship between accuracy and the number of required annotations. In Fig. 3A, we compute the necessary number of annotations required to achieve 80% of classification accuracy with benchmark methods, OpenLabCluster and OpenLabCluster-V on the Home-Cage Mouse dataset. We observe that AL methods require only 15–20% of the annotated segments to achieve 80% of classification accuracy, whereas benchmark methods (KNN, SVM, C, A-SOiD, SimBA) require roughly nine times as many annotations as the optimal AL approaches. Among the AL methods, the MI and CS embedded variants of OpenLabCluster and OpenLabCluster-V achieve 80% accuracy with approximately 400 annotated samples. The variant with the TOP selection method turns out to be slightly less effective, requiring about 700 annotations.

We further visualize the effectiveness of pertaining (C vs. OpenLabCluster) under varying annotation budgets in Fig. 3B. We observe that for most cases, OpenLabCluster methods lead to higher accuracy for a given number of annotations than the counterpart, encoder-only classifier. The curves indicating the accuracy of various OpenLabCluster AL methods (red, green, blue) have a clear gap between them and C curve (darkgray), especially in the mid-range of the number of annotations. However, the improvement of OpenLabCluster over C is less pronounced on the *C. elegans* dataset, likely due to the dataset’s relative simplicity. In Fig. 3C, we further examine class-wise confusion matrices for the Zebrafish dataset on 4 annotation budgets (5%, 10%, 20%, 100%). From visual inspection, it appears that the matrix that corresponds to 20% annotations is close to the matrix that corresponds to 100% annotations. This proximity suggests that the annotation of the full dataset might be redundant. Indeed, further inspection of Fig. 3C, indicates that samples annotated as LCS and BS classes (y-axis) by the unsupervised learning method are likely to be predicted as the LLC (x-axis) by OpenLabCluster. One possibility for the discrepancy could be annotation errors of the prior clustering method, which are taken as the ground truth. Re-examination of the dynamics of some of the features (e.g. tail angle) further supports this hypothesis and demonstrates that the methods in OpenLabCluster can potentially identify the outlier segments whose annotation settles the organization of the Behavior Classification Map (for more details see in Appendix 5.2).

#### Organization of the Latent Representation

Our results indicate that the Latent Representation captured by the OpenLabCluster encoder-decoder and the classifier are able to better organize behavioral segments in comparison to direct embeddings of body keypoints. We quantitatively investigate such an organization with the Monkey dataset, for which ground truth annotations and segmentation are unavailable. Specifically, we obtain the Cluster Map of the segments with OpenLabCluster and then annotate 5% of segments through AL methods which further train the encoder-decoder and the classifier, generating a revised Cluster Map. We then depict the 2D tSNE projection of the Latent Representation and compare it with the 2D tSNE projection of body keypoints in Fig. 4. Indeed, it can be observed that within the Cluster Map, segments are grouped into more distinct and enhanced clusters. To measure the clustering properties of each embedding, we apply clustering metrics of Calinski-Harabasz (CHI) [80] and Davies-Bouldin (DBI) [81]. CHI measures the ratio of inter- and intra-cluster dispersion, with larger values indicating better clustering. DBI measures the ratio of inter-cluster distance to intra-cluster distance, with lower values indicating better clustering. CHI and DBI are shown in the bottom left of Fig. 4 considering the various number of clusters (2, 4, 6, 8, 10, 12). The comparison shows that the CHI index is higher for the Cluster Map than the embedding of the keypoints regardless of the number of clusters being considered and is monotonically increasing with the number of clusters. The DBI index for the Cluster Map is significantly lower than the index of the embedding of the keypoints for up to 10 clusters with the minimum at 6 and 10 clusters. This is consistent with the expectation that clustering quality will be consistent with the number of behavioral types. Indeed in these behaviors, there are 6 major behavioral classes that can be identified and possibly several transitional ones. We show in Fig. 4 (right) the Behavioral Classification Map generated by OpenLabCluster along with frames from representative segments of each class.

## 3 Discussion

In this paper, we introduce OpenLabCluster, a novel toolset for quantitative studies of animal behavior from video recordings in terms of automatic grouping and depiction of behavioral segments into clusters and their association with behavioral classes. OpenLabCluster works with body keypoints which describe the pose of the animal in each frame, and across frames reflecting the kinematic information of the movement that is being exhibited in the segment. The advancement and the availability of automatic tools for markerless pose estimation in recent years allows the employment of such tools in conjunction with OpenLabCluster for performing almost automatic organization and interpretation of a variety of ethological experiments.

The efficacy of OpenLabCluster is attributed to two major components; (i) Unsupervised pre-training process which groups segments with similar movement patterns and disperses segments with dissimilar movement patterns (Clustering); (ii) Automatic selection of samples of segments for association with behavioral classes (AL) through which all segments class labels are associated (classification) and the clustering representation is being refined.

We evaluate OpenLabCluster performance on various datasets of recorded animal species freely behaving, such as Home-Cage Mouse, Zebrafish, *C. elegans* and Monkey datasets. For the datasets for which ground-truth labels have been annotated, we show that OpenLabCluster classification accuracy exceeds the accuracy of a direct deep-classifier for most annotation budget even when all segments in the training set have been annotated. The underlying reason for the efficacy of OpenLabCluster is the unsupervised pre-training stage of the encoder-decoder which establishes similarities and clusters segments with the Latent Representation of the encoder-decoder. Such a representation turns out to be useful in informing which segments could add semantic meaning of the groupings and refine the representation further.

In practice, we observe that even a sparse annotation of a few segments (5%-20% of the training set) chosen with appropriate AL methods would boost clustering and classification significantly. Classification accuracy continues to improve when more annotations are performed, however, we also observe that the increase in accuracy is primarily in the initial annotation steps, which demonstrates the importance of employing clustering in conjunction with AL selection in these critical steps. Indeed, our results demonstrate that among different AL approaches, more direct approaches such as Top, are not as effective as others considering the need to include more metrics quantifying uncertainty and similarity of the segments.

As we describe in the Methodology section, OpenLabCluster includes advanced techniques of unsupervised and semi-supervised neural network training through AL (Appendix 5.1). Inspired by the DeepLabCut project [35, 38, 36], we implement these techniques jointly with a graphic user interface to enable scientists to use the methodology to analyze various ethological experiments with no deep learning technical expertise. In addition, OpenLabCluster is an open-source project and is designed such that further methodologies and extensions would be seamlessly integrated into the project by developers. Beyond ease of use, the graphic interface is an essential part of OpenLabCluster functionality, since it visually informs the scientists of the outcomes in each iteration step. This provides the possibility to inspect the outcomes and assist with additional information “on-the-go”. Specifically, OpenLabCluster allows for pointing at points (segments) in the maps, inspecting their associated videos, adding or excluding segments to be annotated, working with different low-dimensional embeddings (2D or 3D), switching between AL methods, annotating the segments within the same interface, and more.

## 4 Materials and Methods

Existing approaches for behavior classification from keypoints are supervised and require annotation of extensive datasets before training [30, 33]. The requirement limits the generalization of classification from one subject to another, from animal to animal, from a set of keypoints to another, and from one set of behaviors to another due to the need for re-annotation when such variations are introduced.

In contrast, grouping behavioral segments into similarity groups (clustering) typically does not require annotation and could be done by finding an alternative representation of behavioral segments reflecting the differences and the similarities among segments. Both classical and deep-learning approaches address such groupings [4, 48]. Notably, clustering is a ‘weaker’ task than classification since does not provide the semantic association of groups with behavioral classes, however, could be used as a preliminary stage for classification. If leveraged effectively, as a preliminary stage, clustering can direct annotation to minimize the number of segments that need to be annotated and at the same time to boost classification accuracy.

OpenLabCluster, that is primarily based on this concept, first infers a *Cluster Map* and then leverages it for automatic selection of sparse segments for annotation (AL) that will both inform behavior classification and enhance clustering. It iteratively converges to a detailed *Behavior Classification Map* where segments are grouped into similarity classes and each class is homogeneously representing a behavioral state. Below we describe the components.

### Clustering

The inputs into OpenLabCluster denoted as 𝒳, are sets of keypoints coordinates (2D or 3D) or kinematics features for each time segment along with the video footage (image frames that correspond to these keypoints). Effectively, each input segment of keypoints is a matrix with the row dimension indicating the keypoints coordinate, e.g. the first row will indicate the x-coordinate of the first keypoint and the second row will indicate the y-coordinate of the first keypoint and so on.

The first stage of OpenLabCluster is to employ a Recurrent Neural Network (RNN) encoder-decoder architecture that will learn a Latent Representation for the segments as shown in Fig. 5. The encoder is composed of *m* bi-directional gated recurrent units (bi-GRU) [82] sequentially encoding time-series input into a Latent Representation (latent vector in ℝ^*m*^ space). Thus each segment is represented as a point in the Latent Representation ℝ^*m*^ space. The decoder is composed of uni-directional GRUs that receive as input the latent vector and decode (*reconstruct*) the same keypoints from the latent vector. Training optimizes encoder-decoder connectivity weights such that the reconstruction loss, the distance between the segment keypoints reconstructed by the decoder and the input segment, is minimized, see the Appendix for further details (Section 5.1). This process reshapes the latent vector points in the Latent Representation space to better represent the segments similarities and distinctions.

**Figure 5.**
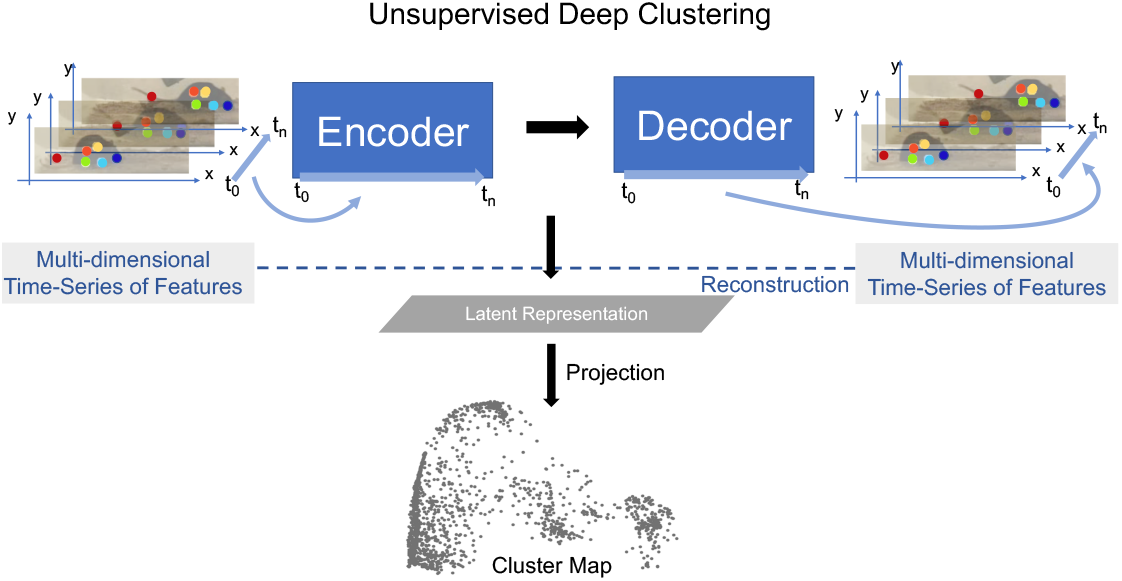
Latent Representation is learned by performing the reconstruction task using an encoder-decoder structure. Latent vectors (last state of the encoder) are projected onto low dimensional space with various dimension reduction techniques to visualize the Latent Representation which constitute the Cluster Map. Mouse images are reproduced from frames of videos in the dataset of [21].

To visualize the relative locations of segments in the Latent Representation, OpenLabCluster implements various dimension reductions (from ℝ^*m*^ → ℝ^2^ or ℝ^*m*^ → ℝ^3^), such as PCA, tSNE, UMAP, to obtain Cluster Maps, see Fig. 5-bottom. Thus each point in the Cluster Map is a reduced-dimensional Latent Representation of an input segment. From inspection of the Cluster Map on multiple examples and benchmarks, it can be observed that the Latent Representation clusters segments that represent similar movements into the same clusters, to a certain extent, typically more effectively than an application of dimension reduction techniques directly to the keypoints segments [60, 83].

### Classification

To classify behavioral segments that have been clustered, we append a classifier, a fully connected network, to the encoder. The training of the classifier is based on segments that have been annotated and minimizes the error between the predicted behavioral states and the behavioral states given by the annotation (cross-entropy loss). When the annotated segments well represent the states and the clusters, the learned knowledge is transferable to other unlabeled segments. AL methods such as Cluster Center (*Top*), Core-Set (*CS*) and Marginal Index (*MI*) aim to select such representative segments by analyzing the Latent Representation. *Top* selects representative segments which are located at the centers of the clusters (obtained by Kmeans [84]) in the Latent Representation space. This approach is effective at the initial stage. *CS* selects samples that cover the remaining samples with minimal distance [85]. *MI* is an uncertainty-based selection method, selecting samples that the network is most uncertain about. See Appendix 5.1 for further details regarding these methods. Once segments for annotation are chosen by the AL method, OpenLabCluster highlights the points in the Cluster Map that represent them and their associated video segments, such that they can be annotated within the graphic interface of OpenLabCluster (choosing the most related behavioral class). When the annotations are set, the full network of encoder-decoder with appended classifier is re-trained to perform classification and predict the labels of all segments. The outcome of this process is the Behavior Classification Map which depicts both the points representing segments in clusters and associated states labels with each point (color) as illustrated in Fig. 6. In this process, each time that a new set of samples is selected for annotation, the parameters of the encoder-decoder and the classifier are being tuned to generate more distinctive clusters and more accurate behavioral states classification. The process of annotation and tuning is repeated, typically until the number of annotations reaches the maximum amount of the annotation budget, or when clustering and classification appear to converge to a steady state.

**Figure 6.**
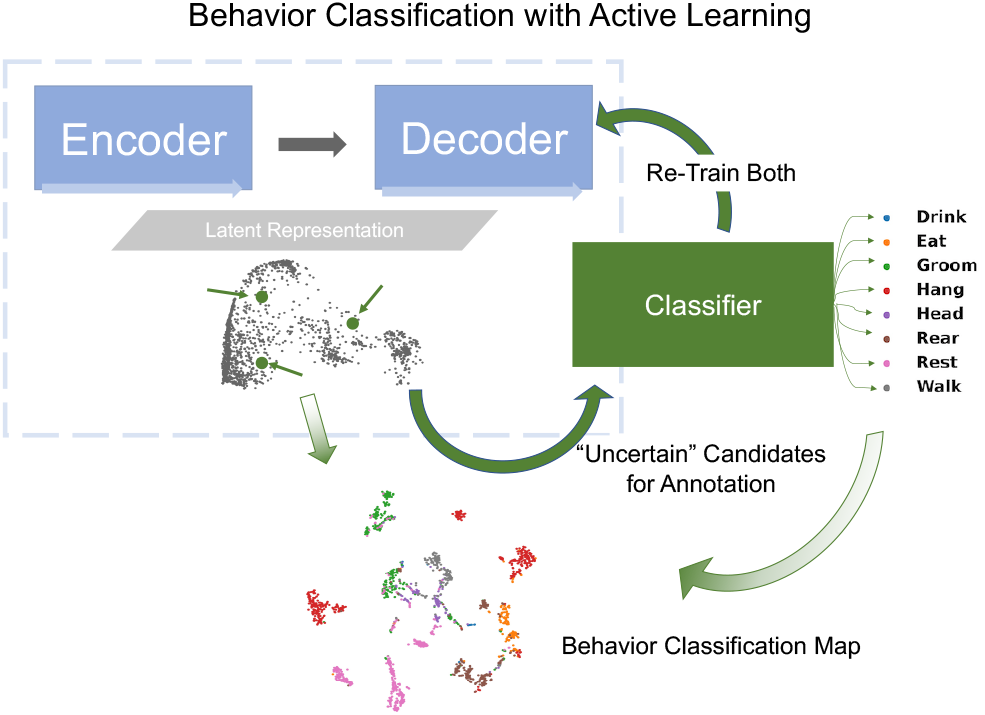
Behavior Classification Map is generated by a fully connected classifier network (green rectangle) which receives the latent vector transformed by the encoder-decoder as input and classifies them into behavior classes (example shown: 8 classes in Home-Cage Mouse Behavior dataset). Behavioral Classification Map is generated from the Cluster Map and indicates the predicted classes of all segments.

### Implementation Details

OpenLabCluster code [86] was developed in University of Washington UW NeuroAI Lab by Jingyuan Li and Moishe Keselman. OpenLabCluster interface is inspired by DeepLabCut [36], which code is used as a backbone for user interface panels, interaction with the back-end, logging, and visualization. OpenLabCluster also uses Google Active Learning Playground code [87] for the implementation of the K-center selection method in the Core-Set AL option. For specific usage please see the third_party folder within the OpenLabCluster code repository [86]. OpenLabCluster is available as a GitHub Repository (OpenLabCluster) https://github.com/shlizee/OpenLabCluster and also can be installed with Package Installer for Python (PIP) *pip install openlabcluster* [88]. The repository includes a manual, instructions, and examples.

### Benchmark Details

As described earlier, OpenLabCluster summarizes keypoints or kinematic features of a temporal segment into a latent representation and then classifies the behavior using this summarized representation. This approach captures the intrinsic dynamics of short behavior prototypes, in contrast to benchmark methods that compute movement features at each timestep via predefined protocols [63, 76] and classify behavior on a per-timestep basis. To ensure fair comparison, we concatenated the frame-wise features within each segment and applied each frame-wise classification method to the resulting representation. Specifically, for KNN [79], SVM [21], and A-SOiD [76], we concatenated the frame-wise features of each action segment and then employed the classifier proposed by each method for behavior recognition. For SimBA [68], movement features were extracted from each frame and integrated with pose-based features. The final representation was formed by concatenating these integrated features across all timesteps within the segment. VAME [61] closely resembles OpenLabCluster by learning a unified representation for entire sequences. In VAME+C, we pre-trained VAME, appended a classifier to its latent feature—which encodes the temporal segment’s dynamics—and fine-tuned the model using a classification loss. For the Home-Cage Mouse, Zebrafish, and *C. elegans* datasets, sequences are pre-segmented so that each segment contains a single behavioral prototype. For the OpenStudio Monkey dataset, which is continuously recorded, we divided the videos into fixed temporal windows. More advanced approaches, such as change-point detection algorithms [89, 90], could also be employed for video segmentation.

## 5 Appendix

### 5.1 Methods

The inputs of OpenLabCluster are multi-dimensional time series representing the coordinates of animal body keypoints or other kinematic features tracked during movement along with video segments frames from which these were extracted. We denote the times-series as 𝒳= {*X*_*u*_ ∪*X*_*l*_}, with *X*_*u*_ representing the sequences in the unlabeled set and *X*_*l*_ in the labeled set, and assume that the segments are unlabeled, 𝒳= *X*_*u*_, in the beginning. A segment **x**_*i*_ ∈χ is represented as a sequence **x**_*i*_ = [*x*_1_, *x*_2_, .., *x*_*t*_, …, *x*_*T*_], where *x*_*t*_ is the vector of features for movement at time *t, x*_*t*_ *∈ R*^*p*^. For the examples we consider here, the number of features is 16, 98, 16, 39 for Home-Cage Mouse, *C. elegans*, Zebrafish and Monkey datasets, respectively. OpenLabCluster includes three main components: (i) An encoder-decoder network that learns to reconstruct sequences and forms Latent Representations. (ii) A classifier network which performs behavior classification. (iii) AL selection which integrates Latent Representation information with the classifier output to optimize classification and annotation.

#### (i) Encoder-Decoder

We adopt the Predict & Cluster encoder-decoder framework which has been shown to achieve self-organizing latent representations [60, 83]. The encoder uses bidirectional Gated Recurrent Units (GRU) and receives **x**_**i**_ ∈ 𝒳 as input. The vector 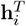 is the latent representation which is the hidden state of the encoder GRU at the last time step *T*. It encodes the dynamic properties of the whole sequence **x**_**i**_ and lies in the latent space *V*, where 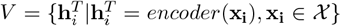, i.e., the space spanned by the latent codes of all sequences. The unidirectional GRU-based decoder receives 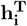 and generates 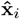 - the reconstruction of the original input sequences. The encoder-decoder network is trained by minimizing the reconstruction loss

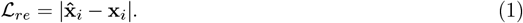

#### (ii) Classification

The classifier is a one-layer fully connected network appended to the encoder. It takes Latent Representation as input and generates the probabilities that a sample belongs to each behavioral states. During training, the classifier is learned to maximize the probability of the annotated behavior state. In other words, with the annotated samples given the classifier output, the classification loss is computed as

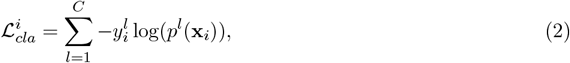

where 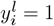 if **x**_*i*_ belongs to class *l*, and 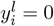otherwise. The complete loss for each sample **x**_*i*_, is then composed from the reconstruction loss 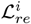 and the classification loss 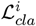 for the annotated samples.

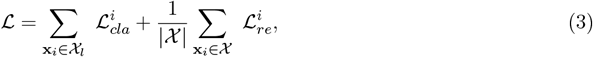

where |𝒳 | is the total number of samples in the dataset. This includes all labeled samples annotated in current and earlier iteration.

#### (iii) AL

There are three AL methods embedded in OpenLabCluster: Cluster Center (Top), Uncertainty with Marginal Index (MI), Core-Set (CS).

**Cluster Center (Top)** leverages clusters information in the latent space to enhance coverage and effectiveness of selected segments (samples). Specifically, *K-Means* clustering is used to transform the latent representation into a collection of clusters 𝒦. The number of clusters *k*

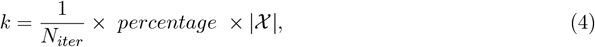

is chosen based on the total number of selection iterations *N*_*iter*_ and the *percentage* of data would be annotated in total. *k* is fixed across selection iterations such that *k* is the number of samples to be annotated and each is located in a different cluster.

**Marginal Index (MI)** is based on the classifier output and measures the difference between the classifier output, evaluating top two difference of *p*. The probability prediction *p* of each class *l* is

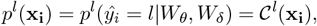

where *p*^*l*^ denotes the probability of a sample to belong to a class *l* among *C* classes (*l* ∈ [1, *C*]) predicted by the classifier 𝒞. 𝒞 indicates the transformation with the classifier. MI is computed as the measure of the confidence, difference between the most probable class and the second most probable class [91, 92], i.e.,

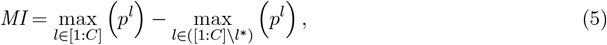

where *l*\* = arg max_*l ∈*[1:*C*]_ (*p*^*l*^).

**Core-Set (CS)** aims to discover a set of samples that can cover unlabeled samples with a certain radius [85]. The algorithm finds a set of samples such that the radius is minimal.

With the labeled samples, the classifier is trained until the classification accuracy on these labeled samples converges. Annotation iteration repeats until reaching the annotation budget.

### 5.2 Zebrafish Behaviors and Dynamics

Zebrafish behavior is subdivided into 13 classes: approach swims (ASs), slow types 1 (S1), slow types 2 (S2), short and long capture swim (SCS and LCS), burst type forward swim (BS), J-turns, high-angle turn (HAT), C-start escape swims (SLC), long latency C-starts (LLC), O-bends, and routine turns (RTs), spot avoidance turn (SAT).

As shown in Fig 3C, OpenLabCluster could “misclassify” samples to belong to LLC class, although they have been annotated by previous analysis as BS or LCS. Since the annotation is obtained by a prior unsupervised learning method and not a case-by-case manual annotation, it could be possible that the annotation is made incorrectly, especially for those segments that are located near the boundaries of clusters. We show several of such conflicting examples in Fig. 7. In the top row, we show the typical dynamics of three ground-truth (GT) classes (BS, LCS, and LLC). Inspecting the prototypical dynamics of the LCS, BS, and LLC, one can find that each profile has unique dynamic profile. For example, LLC profile peaks in the beginning and gradually decays. In the bottom row of Fig. 7, we show three examples of segments classified by OpenLabCluster as *LLC* but marked differently by prior annotation that we consider as ground-truth. It appears that while these examples are similar to LLC profile (having a peak in the beginning and decay) supporting the decision of OpenLabCluster to classify these segments as LLC, the ground truth annotations made by prior clustering analysis are contradictory.

**Figure 7.**
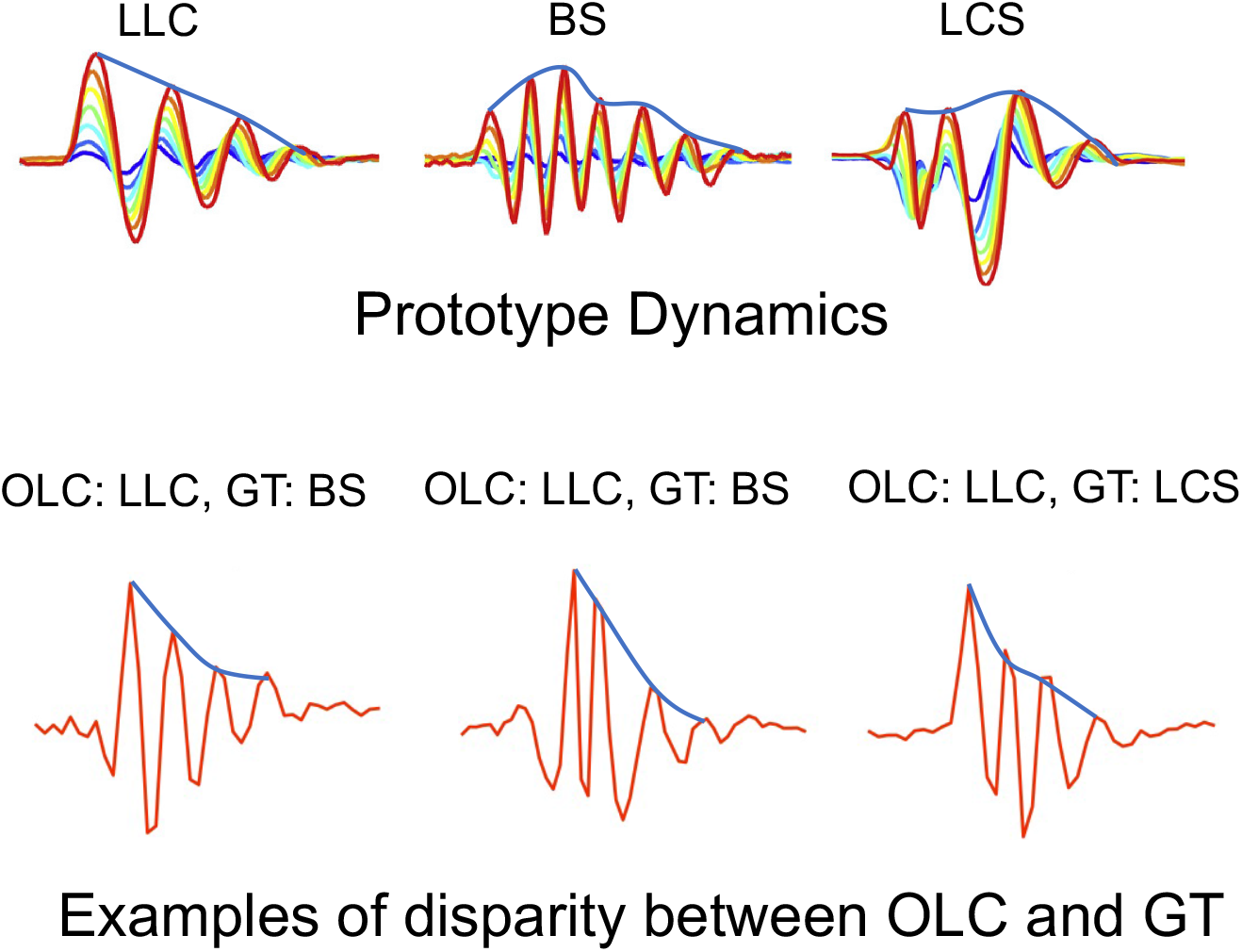
Top: Prototypical dynamics of three behavioral states (LLC, BS, LCS), envelop (blue). Bottom: Three examples of disparity between OpenLabCluster and Ground Truth (GT) classification.

### 5.3 Ablation study

The two components of OpenLabCluster of unsupervised clustering and semi-supervised classification are essential to the accuracy. In Table 3 we examined “ablated” versions of OpenLabCluster to demonstrate the impact of each component; *C* designates a variant that includes a classifier network only. *RC* designates a variant where pretraining phase of OpenLabCluster is ablated, i.e., all the weights of the encoder-decoder and the classifier are randomly initialized. In this scenario, there is no pre-organized Latent Representation. *IRC* designates a variant where AL is not being applied but the encoder-decoder structure as well as the training paradigm including pretraining is kept the same as in OpenLabCluster (with pre-organized Latent Representation). As shown in Table 3, the accuracy of IRC is higher than that of C and RC showing the importance of pre-organized encoder-decoder Latent Representation. OpenLabCluster along with various AL approaches further enhances the accuracy (over IRC) in most cases, especially on the Home-Cage Mouse and the Zebrafish datasets with an average improvement 5.78%. The reason that improvement of OpenLabCluster is not significant on the *C. elegans* dataset could be that the diversity is limited on the dataset, i.e., samples are equally informative for behavior states learning.

**Table 3:** Comparison of OpenLabCluster against its ablated versions: RC, IRC for 5%, 10% and 20% annotations on Mouse, Zebrafish and *C. elegans* datasets.

|  | Mouse |  |  | Zebrafish |  |  | *C. elegans* |  |  |
| --- | --- | --- | --- | --- | --- | --- | --- | --- | --- |
| Label(%) | 5 | 10 | 20 | 5 | 10 | 20 | 5 | 10 | 20 |
| Label(#) | 143 | 286 | 571 | 265 | 530 | 1059 | 27 | 55 | 109 |
| C | 55.2 | 54.1 | 64.5 | 72.5 | 70 | 75.8 | 71.3 | 75.2 | 77.5 |
| RC | 53.7 | 54.1 | 63.9 | 66.3 | 72.3 | 74.8 | 71.0 | 76.4 | 79.1 |
| IRC | 63.0 | 66.6 | 76.1 | 72.8 | 75.9 | 79.0 | 73.8 | **77.0** | 77.3 |
| OpenLabCluster CS | **65.3** | 75.4 | **82.2** | 71.9 | 76.6 | 79.6 | 73.5 | 64.3 | 75.4 |
| OpenLabCluster Top | 58.5 | 58.4 | 76.4 | 72.0 | 77.1 | 80.1 | 76.5 | 76.5 | **77.8** |
| OpenLabCluster MI | 63.4 | **76.3** | 78.9 | **74.2** | **79.1** | **81.1** | **76.7** | 76.8 | 77.0 |

